# Brainstem cholinergic modulation of the thalamocortical activity in urethane anesthetized mice

**DOI:** 10.1101/2021.05.12.443746

**Authors:** Y Audrey Hay

## Abstract

In mammals, sleep consists in the recurrence of two main stages the rapid eye movement (REM) sleep and the slow wave sleep (SWS). The full expression of sleep rhythms requires an intact thalamocortical loop, and its modulation by neuromodulators such as acetylcholine. A high tone of acetylcholine is observed during REM sleep while a low tone of acetylcholine modulates the cortical slow waves during SWS. Brainstem Cholinergic neurons activity correlates with cortical sleep stages but these neurons do not project directly to the cortex. Instead, they could contribute to cortically-recorded sleep stage modulation via a thalamic relay, in particular via the midline thalamic nuclei. Focusing on the brainstem LDTg cholinergic neurons, I investigated how midline thalamic single unit activity and cortical sleep-like stages are modulated during optogenetic-induced activation or silencing of LDTg cholinergic neurons in urethane anesthetized mice. Thalamic neurons were more active during REM-like than SWS-like stages. Bursting activity predominated during SWS-like while tonic firing was prominent during REM-like stage. Optogenetic silencing of the brainstem LDTg cholinergic neurons abolished REM-like stages and reduced tonic spiking of thalamic neurons. Moreover, during SWS-like, silent Down states were prolonged and thalamic tonic spiking during Up states was reduced. Stimulation of the brainstem LDTg cholinergic neurons had a mild effect on thalamic activity even though tonic discharge was increased. Surprisingly, optogenetic stimulation abolished as well REM-like stages maybe through compensatory mechanisms.

## INTRODUCTION

Sleep and wake states are characterized by the amplitude and frequency of the electrical signal recorded at the surface of the brain and by the muscular activity. In mammals, one distinguishes two main stages, slow wave sleep (SWS) and rapid eye movement (REM) sleep. During REM sleep, cortical neurons are depolarized and show little synchronicity, which gives rise to low amplitude oscillations recorded in the cortical local field potential (LFP). During SWS, cortical neuron activity fluctuates synchronously between depolarized, active periods, called Up states, and hyperpolarized, silent periods, called Down states. Cycles of Up and Down states (UDS) are reflected in the LFP as slow oscillations (0.5-4 Hz) of large amplitude (Steriade et al., 1993a). The full expression of SWS and REM sleep requires an intact thalamocortical loop but is also modulated by neuromodulators from subcortical areas, and in particular by cholinergic inputs.

The thalamus consists in relay nuclei that project mainly to primary sensory cortices and non-specific nuclei that project broadly to sensory and associative cortical areas. During SWS, the thalamus paces the slow oscillations of SWS (Steriade et al., 1993b,c,d; David et al., 2013) and generates a faster oscillation called spindle (8-15 Hz; Dempsey and Morison, 1942; Bonjean et al., 2012). During wake, non specific thalamic neurons provide the excitatory drive necessary to sustain attention (Bolkan et al., 2017; Guo et al., 2017; Schmitt et al., 2017). During SWS, burst firing predomines (Hirsch et al., 1983), often associated to specific phases of the cortical oscillations (Urbain et al., 2020), which vary between thalamic nucleus (Sheroziya and Timofieev, 2014; Gent et al., 2018; Varela and Wilson, 2020). The activity of cholinergic neurons is also correlated with arousal states, cholinergic neurons are active during REM sleep and wake and virtually silent during SWS (Boucetta and Jones, 2009), and manipulation of their spiking activity can promote or inhibit arousal states (Steriade et al., 1993e; Van Dort et al., 2015). The thalamocortical loop receives cholinergic inputs from the basal forebrain and the brainstem. Brainstem cholinergic neurons are of particular interest in the regulation of sleep wake cycle as they are part of the historical ascending reticlular activating system (ARAS; Moruzzi and Magoun, 1949). These neurons are located in the laterodorsal tegmental area (LDTg) and pedunculopontine tegmental area (PPTg; Mesulam et al., 1983; Mena-Segovia et al., 2017). LDTg cholinergic neurons only sparsely project to the neocortex (Mesulam et al., 1983; Cornwall et al., 1990) and act on the cortical arousal states through their rostral relays: the non-specific thalamus and the basal forebrain (Steriade et al., 1993e). Here I explore how LDTg neurons modulates the thalamic activity and how this affects cortical LFP.

I first revisited the relation of midline thalamic neurons and cortical LFP during sleep-like stages in the urethane anesthetized mice. By recording simultaneously the cortical LFP and of thalamic single units, I show that urethane anesthesia recapitulates sleep stages and presents regular switches between REM-like and SWS-like activity. The thalamic activity was associated with cortical rhythms, thalamic neurons fired preferentially in burst during SWS-like and tonically during REM-like sleep. Moreover, thalamic neurons fired at specific phases of UDS. Next, coupling electrophysiology and optogenetic manipulation of LDTg cholinergic cells, I investigated how LDTg cholinergic neurons modulates the thalamocortical loop modulation. I show that silencing brainstem cholinergic neurons abolished REM-like sleep on the cortical LFP and reduced thalamic tonic firing. In contrast, activating brainstem cholinergic neurons promoted thalamic tonic firing but had little effect on the cortical LFP signal.

## MATERIALS AND METHODS

### Animals

All experiments were performed in accordance with United Kingdom Home Office regulations (Animals (Scientific Procedures) Act 1986 Amendment Regulations 2012) following ethical review by the University of Cambridge Animal Welfare and Ethical Review Body (AWERB). All animal procedures were authorized under Personal and Project licences held by the author. Mice were group housed in conventional, open cages with food and water ad libitum and maintained on a 12h-12h light-dark cycle, with temperature maintained at 22-24 °C and relative humidity kept at 50-55%. Knock-in homozygous ChAT-CRE mice (Jackson Laboratories, Maine, USA, stock #006410) were crossed with LoxP-ChR2(H134R)-EYFP mice (Ai32, Jackson Laboratories stocks #024109 or #012569) to obtain expression of ChR2 in all cholinergic neurons (ChAT*ChR2) or with RCL-ArchT/EGFP-D mice (Ai40D, Jackson Laboratories stocks #021188) to obtain expression of ArchT in cholinergic cells (ChAT*ArchT).

### Anaesthetised recordings

Mice, aged at least 2 months, were deeply anesthetized via intraperitoneal injections of urethane (1.5 to 1.8 g/kg injected in several shots until loss of reflexes is assessed by paw pinching). While restrained in a stereotaxic apparatus (David Kopf instruments, Phymep, France) mice temperature was monitored and maintained at 34-36 °C using a heating blanket (Basi). A small burr hole was drilled at LDTg coordinates AP = -4.92 mm and ML = 1 mm and an optical fiber (300 μm, 0.29 N.A.; Thorlabs) was descended above the LDTg nuclei at a depth of 3 mm. Light delivery was performed by collimating the optic fibre to a blue laser light (488nm, 15 ± 3 mW at fiber tip; laser 2000) or to a green laser light (532 nm, 10 ± 5 mW; laser 2000). Light delivery was controlled by a shutter whose aperture was driven by custom-made Igor 6.5 procedures. Cortical local field potentials (LFP) were recorded from the prefrontal cortex at coordinates AP = + 1.96 mm, ML = 0.4 mm and DV = 1.2 mm using an extracellular parylene-C insulated tungsten microelectrode (127 μm diameter, 1 MΩ; A-M Systems). Single unit were recorded from the thalamus at coordinates AP = -1.58 mm, ML = 1.3 mm and DV = 3.4 -3.9 mm, using a 20º angle to reach midline thalamic neurons without crossing through the superior sagittal sinus and the ventricles. Single unit recording were performed using glass isolated tungsten electrodes of 5-10 µm tip and 2-10 MOhm impedance (https://www.microelectrodes.net/). Electrical signals were amplified and filtered at 0.1-5 kHz (1800 Microelectrode AC Amplifier; A-M Systems), digitized at 20 kHz using an ITC-18 board (Instrutech, Port Washington, NY) and Igor software (Wavemetrics, Lake Oswego, OR).

### Electrophysiology and Analysis

Cortical LFP signal was low-pass filtered (<120 Hz) and the signal was downsampled from 20 kHz to 1 kHz. Each stimulation procedure was repeated at least 10 times with a minimum of 50 s interval between each trial. LFP was analyzed using custom-made procedures in Igor Pro. LFP traces were subjected to a Morlet wavelet analysis followed by the z-scoring of the power spectrum intensity for each frequency. An instantaneous gamma intensity index was computed by summing up the z-scored intensity in the gamma range (30-80 Hz). REM-like bouts and Up states were identified when the gamma intensity index reached 3-5 SD of the mean gamma intensity index. A aggregation step was performed to remove potential artefacts and aggregate segments of Up and Down states. All Up and Down states were manually assessed before being included in the final analysis and only bouts of well-defined SWS-like were included. REM-like sleep episodes showed comparable gamma intensity index value as Up states and were distinguished from Up states based on their duration (> 3 s), the potential presence of a hippocampus-mediated theta (4-8 Hz) component and a reduced power intensity for low frequency (<4 Hz) initiated in the seconds preceding the REM-like episode.

Unit activity analysis was performed on high-pass filtered signal (300 -3000 Hz). Spikes were detected when their amplitude reached a manually adjusted threshold corresponding to 10-15 SD of the mean signal amplitude. Inter-spike interval plots were computed to discriminate spikes occurring within a burst from those fired tonically. Classically, action potential emitted within 5-20 ms of another spike were classified as belonging to the same burst and action potential emitted at > 30 ms of any other spike being classified as tonic action potential. The temporal relationship between spikes and LFP was evaluated using custom-made procedures in R version 3.4.4 (R Core Team, 2018).

### Statistics

Results are presented as mean ± sem unless otherwise stated. Statistics were performed using R (R Core Team, 2018). Non-parametric Mann-Whitney rank and Kendall rank correlation tests were used throughout the manuscript. Paired tests were performed wherever possible.

## RESULTS

### MTN neurons activity during SWS-like and REM-like

I aimed at describing the dynamics of the thalamocortical loop during sleep and the influence of brainstem cholinergic neurons in shaping the thalamocortical dialogue. During urethane anesthesia, brain activity recapitulates sleep stages, which consists in the spontaneous alternation of SWS-like and REM-like stages. I first asked how the spontaneous firing activity of midline thalamic neurons varies in relation to SWS-like and REM-like stages. Local field potentials (LFP) in the prefrontal cortex and single unit activity in the midline thalamus nuclei were recorded simultaneously (Figure 1A). I compared the activity of 40 thalamic neurons from 21 animals during REM-like and SWS-like sleeps. During SWS-like, the cortical LFP showed low frequency and large amplitude oscillations, while REM-like activity was characterized by low amplitude oscillations, with in some recordings a relative bump in the theta range of frequency (5-10 Hz) probably due to distant field activity from the hippocampus (Figure 1A and 1B). Bouts of SWS-like and REM-like sleep were discriminated based on the slow oscillation power and the duration of active events (see Methods). Thalamic neurons activity was lower during SWS-like compared to REM-like state (SWS-like: 1.53 ± 0.26 Hz; REM-like: 3.50 ± 0.56 Hz; n = 40 neurons; paired Mann-Whitney test, p = 0.006; Figure 1C). However the heterogeneity of firing rates across neurons was high (SWS-like firing range: 0.04-7.56 Hz; REM-like firing range: 0.00-14.05 Hz) and 15 out of 40 neurons increase their activity during SWS-like compared to REM-like sleep (Figure 1C). There was no significant correlation between SWS-like and REM-like firing frequency (Kendall rank correlation coefficient Tau = 0.17; p = 0.13). Thalamic neurons exhibit two modes of firing, phasic discharge of bursts of action potentials fired at 50-300 Hz supported by the voltage-dependent window current T and tonic discharge of isolated action potentials fired at a lower frequency (0.1-30 Hz). Bursting activity and tonic spikes were distinguished by plotting the inter-spike interval for each neuron (Figure 1D). During REM-like sleep, tonic activity prevailed (87.4 ± 2.7 %) while the proportion of spikes fired in bursts was significantly higher during SWS-like sleep (55.4 ± 4.3 %; paired Mann-Whitney test, p = 1.2·10^−10^; Figure 1E). Tonic firing was detected during SWS-like in all 40 neurons while burst firing was absent in 11 out of 40 neurons during REM-like sleep. Consequently, burst frequency was higher during SWS-like compared to REM-like (SWS-like: 0.83 ± 0.15 Hz; REM-like: 0.46 ± 0.11 Hz; paired Mann-Whitney test, p = 0.002; Figure 1F), while tonic firing frequency was higher during REM-like than during SWS-like (SWS-like: 0.70 ± 0.16 Hz; REM-like: 3.04 ± 0.49 Hz; paired Mann-Whitney test, p = 6.5·10^−7^; Figure 1G). Last, there was no significant difference between REM-like and SWS-like in the number of spikes emitted within a burst (SWS-like: 2.49 ± 0.08 spikes / burst; REM-like: 2.47 ± 0.15 spikes / burst; ; paired Mann-Whitney test, p = 0.20). Hence, these results demonstrate that midline thalamic activity is correlated with sleep-like stages.

**Caption Figure 1:**
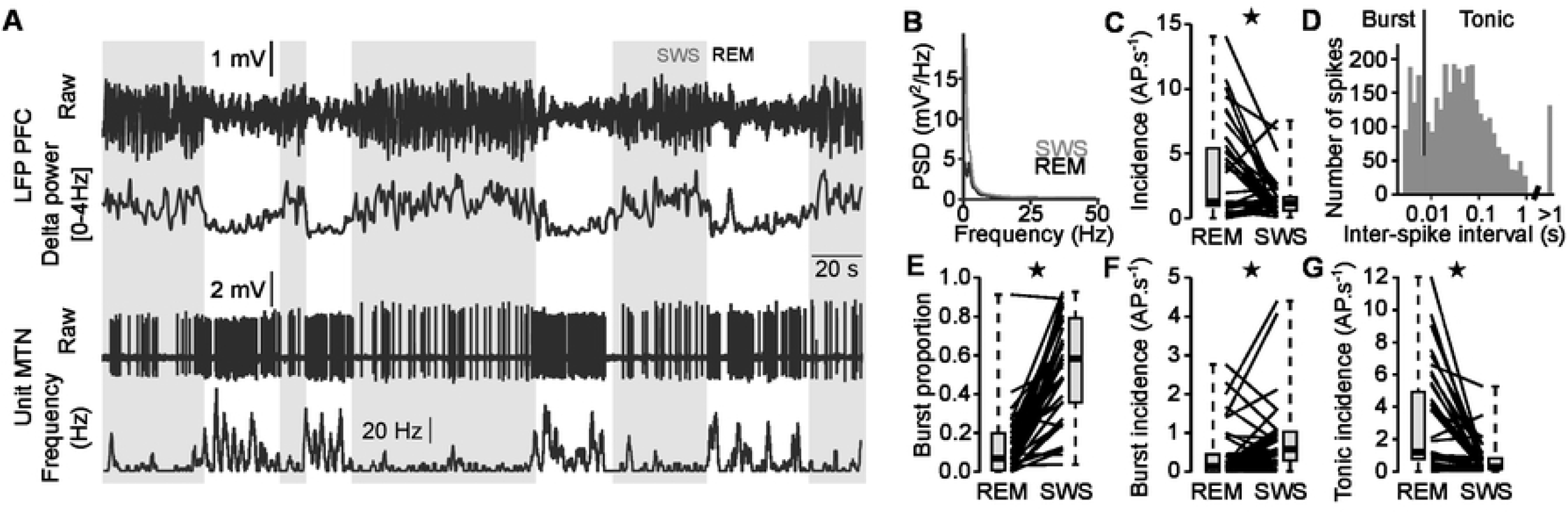
MTN neurons firing activity during urethane-induced REM-like and SWS-like states. **A:** Simultaneous recordings of PFC LFP and a MTN unit exemplifying the higher spiking frequency of MTN neurons during REM-like state compared to SWS-like state. SWS-like is characterized by high delta power (arbitrary unit). **B:** Power spectral density for REM-like state (black) and SWS-like state (grey). **C:** Quantification of the average spiking frequency for each neuron (n = 40) in REM-like and SWS-like states. **D:** Inter-spike interval plot allows for discriminating spikes fired in a burst from tonic action potentials for each individual neurons. **E:** Proportion of spikes fired in a burst in REM-like and SWS-like sleep showing that burst predominates during SWS-like while tonic discharge predominates during REM-like sleep. **F:** Burst firing frequency is higher during SWS-like sleep compared to REM-like sleep. **G:** Tonic firing frequency is higher during REM-like sleep compared to SWS-like sleep.

Next I investigated the thalamic spiking relative to the phase of the UDS. UDS were identified based on cortical multi unit activity, gamma power intensity and slow oscillation waveform (see Methods and Figure 2A). An UDS cycle lasted 1.38 ± 0.08 s and Up and Down states were of similar duration (Up: 0.80 ± 0.03 s; Down: 0.79 ± 0.05 s; n = 21 animals; paired Mann-Whitney test, p = 0.68; Figure 2B); 66.3 ± 2.6 % of the spikes were fired during Up states (incidence Up states : 2.23 ± 0.21 Hz; incidence Down states: 1.54 ± 0.22 Hz; n = 46; paired Mann-Whitney test, p = 7.10^−4^; Figure 2C), indicating that thalamic neurons fire preferentially during the active phase of the cortical UDS. An additional 6 neurons were analyzed in this sample that corresponds to recordings for which too little REM-like sleep was observed to be quantified. In this sample, 46.9 ± 4.7 % spikes were fired tonically and 53.3 ± 4.2 % spikes were fired within a burst (n = 46 neurons). More specifically, 63.3 ± 2.8 % of tonic spikes occurred during Up states (Up: 0.95 ± 0.14 Hz; Down: 0.70 ± 0.12; n = 46; paired Mann-Whitney test, p = 0.007) and 66.4 ± 3.3 % of bursts occurred during Up states (Up: 1.26 ± 0.16 Hz; Down: 0.99 ± 0.18; n = 46; paired Mann-Whitney test, p = 0.02; Figure 2D). Thus thalamic neurons preferentially fire during Up states, with no specific difference between tonic and bursting activity.

**Caption Figure 2:**
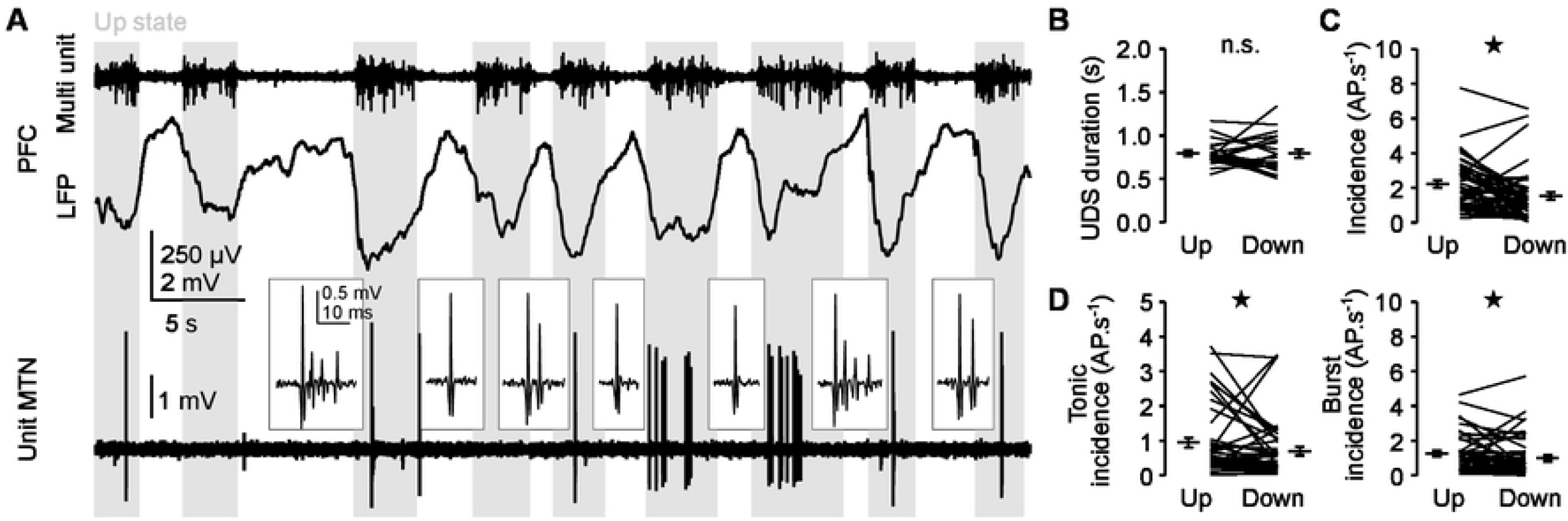
MTN neurons firing activity during cortical Up and Down states of SWS-like. **A:** Simultaneous recordings of PFC LFP, multi-unit activity and MTN unit showing that thalamic neurons fire preferentially during the Up state (grey shadow). Inserts zoom on MTN spikes exemplifying burst and tonic action potential firings. **B:** In urethane-anesthetized mice, Up and Down states are of comparable duration (n = 21 mice). **C-E:** Incidence of spiking activity is higher during Up states compared to Down states for all spikes (**C**), and when tonic (**D**) and burst (**E**) firing are distinguished (n = 46 neurons).

### Silencing LDTg ACh neurons reduces REM-like states and Up state duration and decreases thalamic tonic firing activity

LDTg nucleus is believed to be part of the REM sleep promoting network affecting the cortical rhythms through a thalamic and/or a forebrain relay. I asked how LDTg cholinergic inputs affect the firing activity of the midline thalamic neurons and the cortical LFP. ChAT-CRE mice were crossed with mice harbouring a LoxP-flanked archaerhodopsin-T gene (ArchT), which allows for the selective expression of the neuronal silencer ArchT in cholinergic neurons, and delivered green light (530 nm, 15 ± 3 mW) to the LDTg through an optic fiber implanted above the area. Silencing cholinergic neurons for 30 s abolished REM-like sleep in 4 out of 6 animals and reduced the mean occurrence of REM-like bouts from 2.7 ± 0.6 bouts/100 s to 0.2 ± 0.2 bouts/100 s (n = 6 mice; paired Mann-Whitney test, p = 0.03; Figure 3A-B). SWS-like slow oscillations were also affected by LDTg silencing. UDS incidence decreased (Ctrl incidence: 0.54 ± 0.04 Hz; ArchT incidence: 0.42 ± 0.02 Hz; n = 6 mice; paired Mann-Whitney test, p = 0.03; Figure 3C-D), Down state duration increased (Ctrl Down duration: 0.88 ± 0.11 s; ArchT Down duration: 1.33 ± 0.23 s; n = 6; paired Mann-Whitney test, p = 0.03; Figure 3E) and Up state duration was decreased (Ctrl Up duration: 0.77 ± 0.01 s; ArchT Up duration: 0.64 ± 0.03 s; n = 6 mice; paired Mann-Whitney test, p = 0.03; Figure 3F). Thus silencing LDTg cholinergic neurons affect both REM-like and SWS-like stages by decreasing the time spent in active states.

**Caption Figure 3:**
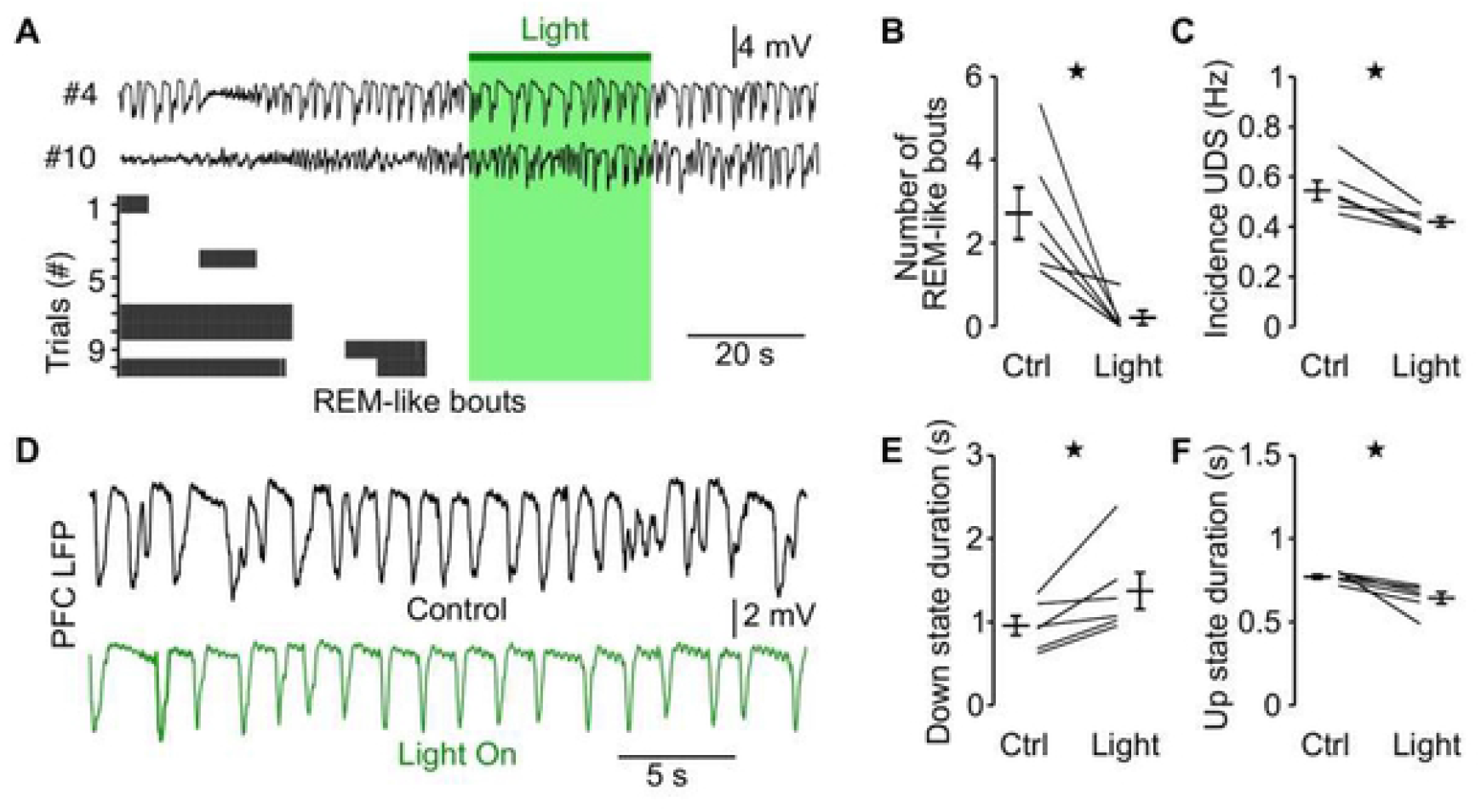
Inhibition of LDTg ACh neurons abolishes REM-like sleep and decreases Up state duration. **A-B:** Silencing of LDTg ACh neurons abolishes REM-like bouts. **A:** Example traces in control condition and during silencing of LDTg neurons (green shade) of PFC LFP. Bouts of REM-like sleep (grey) were absent during silencing. **B:** Quantification of number of REM-like bouts recorded in the PFC across 10 repeats of control and silencing of ACh neurons. **C-F:** Silencing of LDTg ACh neurons reduces time in active state during UDS. **C:** Example traces of PFC LFP shows shorter Up states and longer Down states during LDTg ACh neurons silencing. **D:** Incidence of UDS is reduced, **E:** Down state duration is increased, and **F:** Up state duration is reduced (n = 6 mice).

We investigated whether thalamic neurons firing was affected by LDTg silencing and thus to which extent the cortical altered LFP could be explained by an alteration of the thalamic activity. Because no REM-like sleep bouts were observed in 4 out of 6 neurons, we could not analyzed how LDTg silencing affect thalamic neurons firing in REM-like states and focused on the firing during SWS-like (Figure 4A). Frequency of action potential discharge across UDS was decreased in 8 out of 12 neurons when LDTg ACh neurons were silenced but taking all neurons together this decrease was not significant (Ctrl: 1.83 ± 0.58 Hz; ArchT: 1.11 ± 0.23 Hz; n = 12 neurons; paired Mann-Whitney test, p = 0.09; Figure 4A-B). This trend for a decrease of frequency was not specific for action potentials emitted during neither Up nor Down state, which both showed the same trend (Ctrl Up: 2.17 ± 0.63 Hz; ArchT Up: 1.30 ± 0.35 Hz; n = 12 neurons; paired Mann-Whitney test, p = 0.11; Ctrl Down: 1.59 ± 0.56 Hz; ArchT Down: 1.01 ± 0.33 Hz; n = 12 neurons; paired Mann-Whitney test, p = 0.11; Figure 4B). I next distinguished burst and tonic firing. Burst firing frequency was not affected by the inhibition of LDTg ACh neurons (Ctrl freq: 0.43 ± 0.14 Hz; ArchT freq: 0.47 ± 0.14 Hz; n = 12 neurons; paired Mann-Whitney test, p = 0.76; Figure 4C), the number of spike per burst was also not affected (Ctrl: 2.33 ± 0.08 spikes/burst; ArchT: 2.35 ± 0.07 spikes/burst; n = 12 neurons; paired Mann-Whitney test, p = 1) and the distribution in Up and Down states did not change (Ctrl Up: 0.52 ± 0.15 Hz; ArchT Up: 0.56 ± 0.19 Hz; n = 12 neurons; paired Mann-Whitney test, p = 0.84; Ctrl Down: 0.36 ± 0.14 Hz; ArchT Down: 0.41 ± 0.25 Hz; n = 12 neurons; paired Mann-Whitney test, p = 0.84; Figure 4C). Conversely, a reduction of tonic firing frequency was observed when LDTg ACh neurons were inhibited (Ctrl freq: 1.36 ± 0.55 Hz; ArchT freq: 0.65 ± 0.27 Hz; n = 12 neurons; paired Mann-Whitney test, p = 0.04; Figure 4D). This change was not specific for Up or Down phase of the cycle (Ctrl Up: 1.64 ± 0.60 Hz; ArchT Up: 0.76 ± 0.29 Hz; n = 12 neurons; paired Mann-Whitney test, p = 0.03; Ctrl Down: 1.23 ± 0.52 Hz; ArchT Down: 0.61 ± 0.26 Hz; n = 12 neurons; paired Mann-Whitney test, p = 0.046; Figure 4D). Thus, silencing LDTg cholinergic neurons abolished REM-like sleep and the active phase of the SWS-like, which was accompanied by a reduction of thalamic tonic firing. These results suggest that the thalamus could relay to the neocortex the LDTg inputs and contribute to the active states through tonic firing.

**Caption Figure 4:**
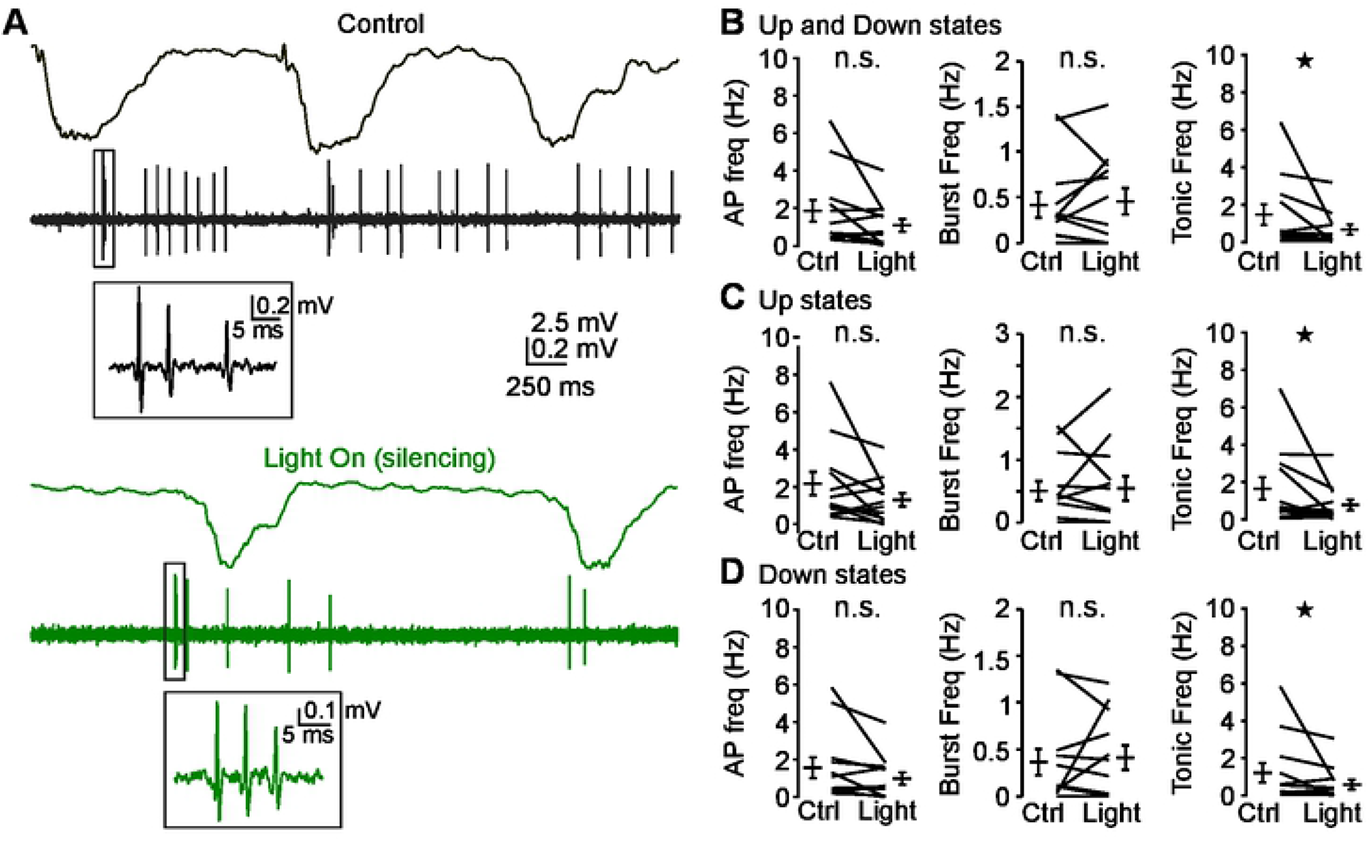
Inhibition of LDTg ACh reduces thalamic tonic firing during SWS-like. **A:** Example of simultaneous recordings from cortical LFP and MTN single unit in control condition (black traces) and during LDTg silencing (green traces). **B-D:** Quantification of the incidence of action potential discharge of thalamic neurons across Up and Down states, taking into account all spikes (left column), burst firing only (middle column) and tonic firing only (right column) during the whole Up and Down state cycle (**B**), Up state only (**C**) and Down state only (**D**). This analysis reveals a specific reduction of tonic firing at all stages of the Up and Down state cycle (n = 12 neurons).

### Stimulation of LDTg ACh neurons does not affect UDS but increases thalamic tonic firing activity

I last investigated how stimulation of LDTg affects sleep-like stages and thalamic firing activity. ChAT-CRE mice were crossed with LoxP-flanked channelrhodopsin-2 mice (ChR2) to selectively express the neuronal activator ChR2 in cholinergic neurons. Blue light (488 nm, 10 ± 2 mW) was delivered to the LDTg through an optic fiber implanted above the area. I recorded 15 neurons from 8 ChAT*Ai32 mice and performed 30-s long blue light pulses repeated every 2 to 5 minutes (Figure 5 and 6). I first looked at REM-like and SWS-like activity (Figure 5A). Unexpectedly, the occurrence of REM-like bouts was strongly reduced during stimulation (Ctrl: 4.0 ± 0.4 bouts / 100 s; ChR2: 1.0 ± 0.4; n = 8 animals; paired Mann-Whitney test, p = 0.01) while Up and Down states incidence or duration were not affected by the stimulation (Ctrl incidence UDS: 0.56 ± 0.03 Hz; ChR2 incidence UDS: 0.57 ± 0.02 Hz; paired Mann-Whitney test, p = 0.29; Ctrl Up duration: 0.86 ± 0.05 s; ChR2 Up duration: 0.92 ± 0.06 s; paired Mann-Whitney test, p = 0.20; Ctrl Down duration: 0.83 ± 0.06 s; ChR2 Down duration: 0.82 ± 0.05 s; n = 8; paired Mann-Whitney test, p = 0.84). Thus LDTg stimulation in the anesthetized mouse has little influence on SWS-like sleep but reduces the occurrence of REM-like sleep during the stimulation.

**Caption Figure 5:**
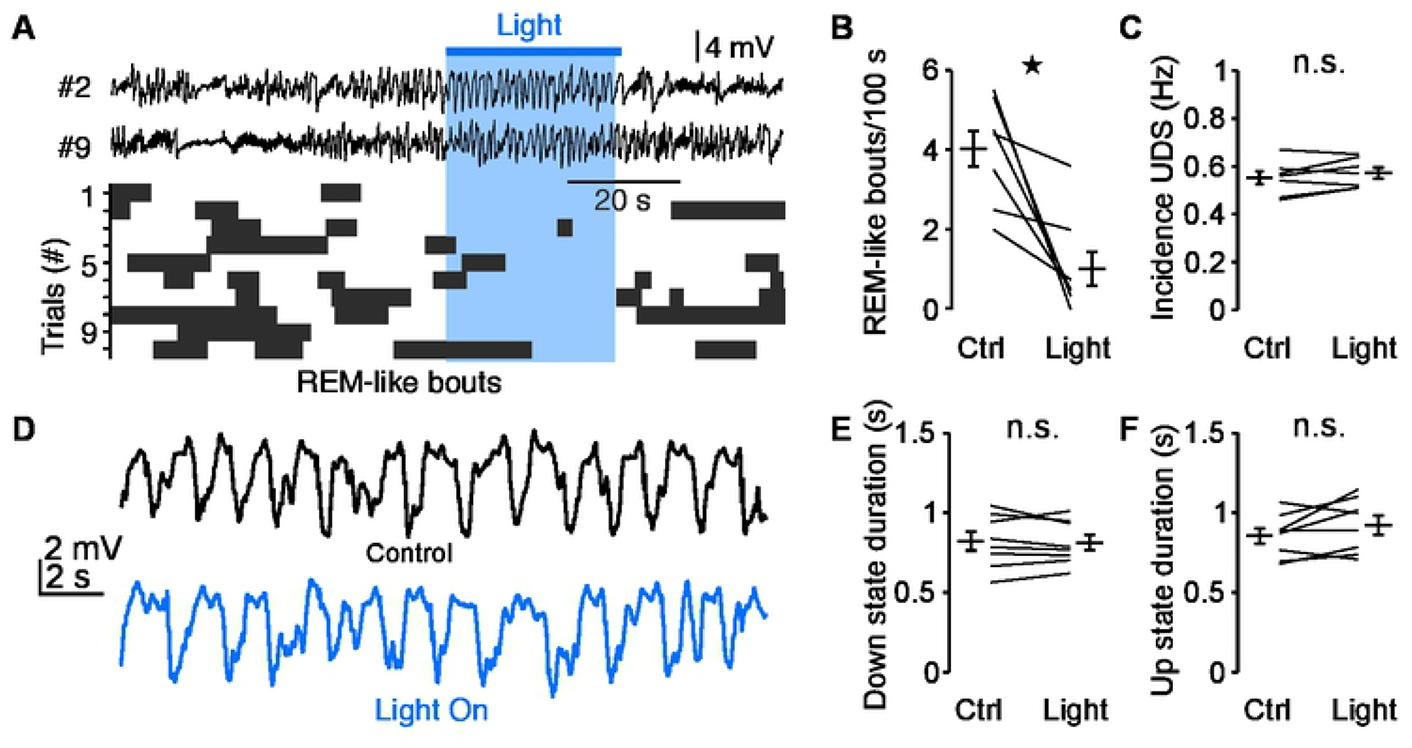
Stimulation of LDTg ACh neurons does not affect Up and Down state duration in the PFC. **A-B:** Excitation of LDTg ACh neurons reduces REM-like bouts incidence. **A:** Example traces of PFC LFP in control condition and during light stimulation of LDTg neurons (blue shade). Incidence of REM-like sleep bouts (dark grey) is reduced during light stimulation. **B:** Quantification of number of REM-like bouts recorded in the PFC across 10 repeats of control and light stimulation of ACh neurons. **C-E:** Light stimulation of LDTg ACh neurons does not affect Up and Down state cycle. **C:** Incidence of UDS is not affected by light stimulation. **D:** Example traces of PFC LFP shows no effect on Up and Down states morphology during LDTg ACh neurons excitation (blue trace) compared to control conditions (black trace). Down (**E**) and Up (**F**) states durations are not affected by light stimulation (n = 8 mice).

**Caption Figure 6:**
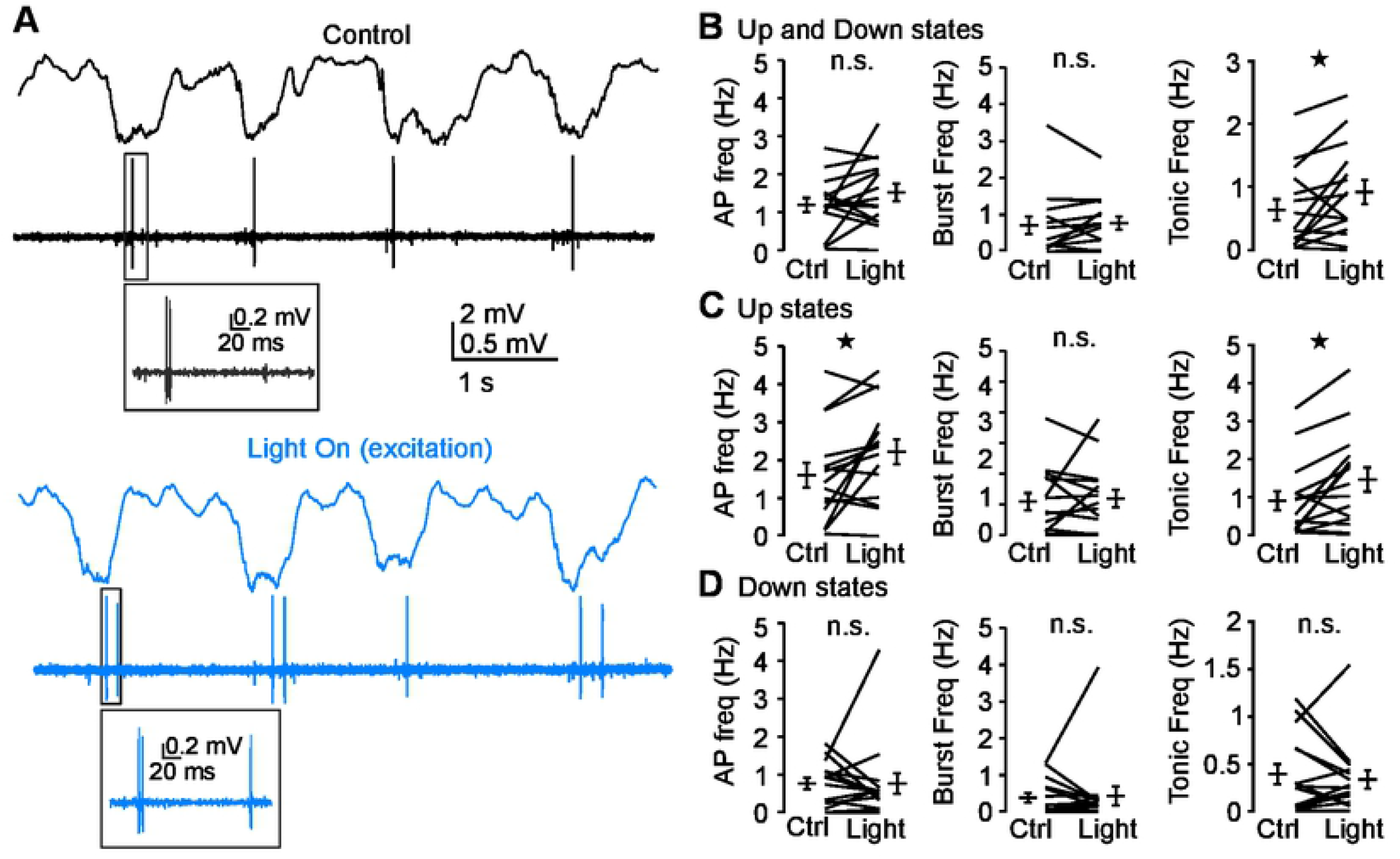
Stimulation of LDTg ACh neurons increases MTN tonic firing specifically during Up states. **A-B:** Example of simultaneous recordings from cortical LFP and MTN single unit in control (black traces) and in light on (blue traces) conditions. **B-D:** Quantification of the incidence of action potential discharge of thalamic neurons across Up and Down states, taking into account all spikes (left column), burst firing (middle column) and tonic firing (right column) separately during the whole Up and Down state cycle (**B**), Up state only (**C**) and Down state only (**D**). This analysis reveals a specific increase of tonic firing during Up states (n = 15 neurons).

Likewise in the MTN, during SWS-like stage, light stimulation did not alter the mean frequency of action potential discharge (Ctrl: 1.19 ± 0.19 Hz; ChR2: 1.52 ± 0.23 Hz; n = 15 neurons; paired Mann-Whitney test, p = 0.28; Figure 6A and 6B). However, looking specifically at spikes emitted during Up or Down states, a significant increase of action potential discharge was observed during Up states (Ctrl Up: 1.60 ± 0.32 Hz; ChR2 Up: 2.22 ± 0.33 Hz; paired Mann-Whitney test, p = 0.035; Figure 6D) but not during Down states (Ctrl Down: 0.78 ± 0.15 Hz; ChR2 Down: 0.78 ± 0.27 Hz; paired Mann-Whitney test, p = 0.53). Looking separately at burst and tonic firing, I observed an increased of tonic firing frequency (Ctrl: 0.64 ± 0.17 Hz; ChR2: 0.92 ± 0.19 Hz; n = 15 neurons; paired Mann-Whitney test, p = 0.04; Figure 6B) and an increase of number of tonic spike per UDS cycle (Ctrl: 1.09 ± 0.29 tonic spike per cycle; ChR2: 1.60 ± 0.36 burst per cycle; n = 15 neurons; paired Mann-Whitney test, p = 0.03). Increase of tonic firing was specific to the Up states phase (Ctrl Up: 0.89 ± 0.26 Hz; ChR2 Up: 1.45 ± 0.32 Hz; n = 15; paired Mann-Whitney test, p = 0.015), while no change of tonic spike frequency was observed during Down states (Ctrl Down: 0.39 ± 0.11 Hz; ChR2 Down: 0.33 ± 0.10 Hz; n = 15; paired Mann-Whitney test, p = 0.49). Conversely, neither the burst frequency (Ctrl: 0.55 ± 0.14 Hz; ChR2: 0.59 ± 0.14 Hz; n = 15 neurons; paired Mann-Whitney test, p = 0.75; Figure 6B) nor the number of burst per UDS cycle (Ctrl: 0.85 ± 0.20 burst per cycle; ChR2: 0.93 ± 0.21 burst per cycle; n = 15 neurons; paired Mann-Whitney test, p = 0.66) were affected by optogenetic stimulation of the LDTg cholinergic neurons. Burst firing was not redistributed across UDS by light stimulation (Ctrl Up: 0.71 ± 0.24 Hz; ChR2 Up: 0.77 ± 0.19 Hz; n = 15; paired Mann-Whitney test, p = 0.40; Ctrl Down: 0.39 ± 0.12 Hz; ChR2 Down: 0.44 ± 0.25 Hz; n = 15; paired Mann-Whitney test, p = 0.84; Figure 6C-D) and the number of spikes fired in a burst was unchanged (Ctrl: 2.43 ± 0.12 spikes/burst; ChR2: 2.43 ± 0.11 spikes/burst; n = 12 neurons; paired Mann-Whitney test, p = 1).

Thus, LDTg ACh neurons stimulation increased tonic firing but not burst firing, and tonic firing increased specifically during Up states, but this increase of thalamic activity had no directly measurable impact on the cortical LFP.

## DISCUSSION

Here I examined the relation between the MTN firing activity and cortical LFP during sleep-like stages in the anesthetized mouse and its modulation by brainstem cholinergic inputs. The main results are: 1) MTN neurons fire primarily in burst during SWS-like and tonically during REM-like sleep; 2) MTN neurons are more active during the Up state of the UDS; 3) silencing of cholinergic neurons abolishes REM-like stages, decreases Up state duration and reduces MTN tonic firing rate; 4) in the urethane anesthetized mouse cholinergic neuron stimulation does not affect the cortical LFP but promotes MTN tonic firing.

### MTN activity across cortical states of vigilance

The results presented here demonstrate that midline thalamic neurons activity is correlated with the cortical sleep-like states in the urethane anesthetized mouse (Figure 1 and 2). As reported during natural sleep, midline thalamic neurons were in average more active during REM sleep than SWS (Gent et al., 2018; Hauer et al., 2019). However, the difference of firing rate between SWS-like and REM-like sleep (twofold in this study, Figure 1) was not as marked as during natural sleep, when the mean activity was up to five times higher during REM sleep (Honjoh et al., 2015; Gent et al., 2018). This difference might be due to the fact that, in my sample, one third of the recorded neurons decreased their activity during REM-like stage. Because thalamic neurons are more active during REM sleep (Honjoh et al., 2015; Gent et al., 2018), the presence of silenced neurons in this sample was surprising. How a neuron would fire during REM-like state was not predictable from its activity during SWS-like sleep (Figure 1), nor from its location in the MTN. Three main reasons then can account for the presently reported silenced neurons: 1) how neurons were sampled, 2) the difference between natural REM sleep and urethane anesthesia REM-like sleep, and 3) the sensitivity of midline thalamic neurons to neuromodulators. 1) In this report, the mean firing rate was 5 times lower for both REM-like and SWS-like state than reported in naturally sleeping mice (Gent et al., 2018). However, the firing rate here was similar as in other anesthetized studies (Zimmerman and Grace, 2018) and neurons with very low baseline activity during sleep have been reported in sensory thalamic nuclei (Urbain et al., 2019). So despite the fact that SWS-like firing rate was not predictable of the REM-like firing rate, it is possible that neurons with low activity were missed in some natural sleep studies. 2) On the other hand, there is little doubt that urethane anesthesia, despite being the most accurate anesthetics at recapitulating sleep-like stages (Pagliardini et al., 2012, 2013), presents different features from natural sleep. In particular, the neuromodulatory state might be different and one can hypothesized that the heterogeneity reported here arises from different sensitivity to neuromodulators and/or varying modulatory state across recordings. Variable modulatory state is however unlikely as each neuron was recorded for at least 30 minutes to sample several REM-like episodes and neurons showed a consistent behavior across REM episodes. Thus it unlikely that a fluctuating neuromodulatory state explains the presence of neurons silent during REM in this study. Moreover, neurons active and silent during sleep were recorded from the same mouse ruling out inter-animal variability. Thus one might speculate that these neurons are a feature of MTN, maybe due to their differential sensitivity to neuromodulators, but this would need to be confirmed in natural sleep.

### Preventing sleep stage transition with brainstem cholinergic neurons silencing

Cholinergic neurons from the LDTg and PPTg are part of the historical reticular arousal system whose stimulation promotes arousal (Moruzzi and Magoun, 1949; Steriade et al., 1993e). LDTg/PPTg cholinergic neurons firing rate is high during REM(-like) and awake state and low during SWS(-like; Menia-Segovia et al., 2008; Boucetta et al., 2014; Cissé et al., 2018) and these neurons show an increase of activity that precedes SWS to REM/wake transition (Petzold et al., 2015). This has suggested a causal role of brainstem cholinergic neurons in triggering SWS to REM stage transition. However, results from experiments activating optogenetically or chemogenetically the PPTg/LDTg cholinergic neurons diverge and the causal role of PPTg/LDTg in promoting SWS to REM transition is still debated (Van Dort et al. 2015; Grace et al., 2014; Kroeger et al., 2017; Cissé et al., 2018). The results presented here, in the urethane anesthetized mouse, indicate that silencing LDTg cholinergic neurons prevent cortical REM-like state (Figure 3), arguing in favour of a necessary role for LDTg activation to permit the transition. In contrast, activating LDTg ACh neurons did not promote REM-like transition (Figure 5), which is against the hypothesis that LDTg ACh neurons activation is sufficient to promote REM-like sleep. However, these results need to be taken very carefully as urethane anesthesia is likely to affect cholinergic modulation as discussed in the next section.

Interestingly, we also show that silencing cholinergic neurons shortens Up state and increases Down state duration (Figure 3). These results are consistent with the observation that cholinergic tone is not null during SWS and low dose of acetylcholine in the cortex can modulate Up and Down state duration (Marrosu et al., 1995; Carracedo et al., 2013; Wester et al., 2013; Lörincz et al., 2015; Hay et al., 2021). However in these studies, acetylcholine was locally applied to the cortex, while here, the cholinergic nucleus targeted only weakly projects to the cortex and only to the caudal area. This suggests that an intermediate structure was involved in modulating the cortical activity.

### Mild influence of optogenetic activation of LDTg neurons on thalamocortical loop

In this study, the optogenetic activation of cholinergic neurons had a limited effect on both LFP and thalamic firing (Figure 5 and 6). This was surprising and in stark contrast with a study in natural sleep, that reports SWS to REM or SWS to awake transition upon light stimulation of brainstem cholinergic neurons (Van Dort et al., 2015). The difference could be due to the milder stimulation protocol used here and in accordance to this hypothesis moderate influence on slow oscillations had been reported under moderate activation of PPTg in urethane animals (Mena-Segovia et al., 2008). Several alternative reasons can account for the mild response reported in this study, 1) the pattern of stimulation, 2) the duration of stimulation, 3) the stimulation of LDTg only and 4) the baseline firing rate of LDTg neurons in urethane anesthesia. 1) Continuous blue light was applied for 30 s long bouts which could have triggered a depolarising plateau of cholinergic neurons membrane potential instead of action potential discharge (Mattis et al., 2012). A depolarizing plateau would probably have led to no acetylcholine release and thus would be of similar effect as the cholinergic neuron silencing. This might explain why we observe no REM-like episode or prolonged Up states during light stimulation. However silencing was associated with an increase of Down states duration and a shortening of Up states (Figure 3), which we did not observe when in the excitation protocol (Figure 5). More importantly optogenetic stimulation was associated with an increase of MTN tonic firing while we observed a reduction of tonic firing upon silencing (Figures 4 and 6). Last, continuous pulse has been shown to promote sustained firing of LDTg neurons (Cissé et al., 2018). Thus it is unlikely that the continuous stimulation protocol had resulted in the silencing of the cholinergic neurons. 2) The protocol used here consisted of relatively short stimulation protocol (30 s) that might have been too short to trigger a SWS-like to REM-like transition. Indeed, Van Dort and collaborators (2015) performed longer stimulation (60 s long minimum) and observed an increase of transition probability peaking roughly at one minute after the onset of the stimulation. Here, REM-like bouts were observed sometimes 60-100 s following the stimulation protocol (Figure 5). More investigations would be needed to determine whether this was coincidental or causal. 3) Alternatively the absence of transition could be explain by the fact that I stimulated the LDTg and not the PPTg. Indeed in natural sleep, the probability to induce a SWS to REM transition was higher upon stimulation of PPTg than LDTg (Van Dort et al., 2015). Here I investigated the role of the MTN as a relay of brainstem cholinergic inputs and thus focused on LDTg that is the primary source of acetylcholine to the MTN (Huerta-Ocampo et al., 2020), however stimulation of PPTg yielded to similarly low effect on cortical LFP (unpublished observation). Thus it is unlikely that the stimulation of LDTg only is the main reason for the mild response observed here. Last, 4) it is possible that in urethane anesthesia LDTg ACh neurons fire at their near maximal rate, which would occlude any effect of the light stimulation. Urethane anesthesia recapitulates sleep like stage (Pagliardini et al., 2012, 2013) as assessed by brain activity recorded from the rostral areas of the brain, however brainstem structures show patterns of activity that are not typical of sleep stages. In particular neurons from Locus coeruleus and from brainstem cholinergic nuclei show rates of activity more closely related to awake states than sleep stages (Mena-Segovia et al., 2008; Boucetta and Jones, 2009; Eschenko et al., 2012) and blockade of cholinergic receptors in the thalamus of anesthetized animals hyperpolarized thalamic neurons indicating of a persistent tone of ACh in the thalamus (Curró Dossi et al., 1991). Thus the already high level activity of the LDTg neurons might have occluded the effect of the optogenetic stimulation. Moreover, brainstem cholinergic neurons are thought to be REM inducing when the brain is in measure to operate the transition. This can be prevented by refractory period or other factors such as closeness to state transition. It is therefore possible that in urethane anesthesia the brain in not in the state that would allow LDTg cholinergic neurons activation to promote a SWS to REM transition.

Recently the views on brainstem cholinergic impact on sleep/wake stages transition has been challenged; despite the fact that stimulation of cholinergic neurons triggers a transient increase of cortical gamma oscillation (Furman et al., 2015; Cissé et al., 2018; Mena-Segovia and Bolam, 2017), some authors suggest that the importance of brainstem cholinergic neurons to promote wakefulness could have been overestimated (Mena-Segovia and Bolam, 2017). The results presented here are more in line with this view and would support the hypothesis that brainstem cholinergic neurons finely tune sleep states rather than promote state transition.

### MTN relays partially brainstem cholinergic inputs to the neocortex through tonic spikes

The thalamus is, with the basal forebrain, the main recipient of brainstem cholinergic fibers in the forebrain, and more specifically the LDTg projects to the MTN (Hallanger et al., 1987; Kitt et al., 1994; Mena-Segovia and Bolam, 2017; Huerta-Ocampo et al., 2020). Non-specific thalamic nuclei receive twice as many cholinergic fibers as the relay nuclei and are specifically sensitive to neuromodulation (Parent and Descarries, 2008; Bayer et al., 2002; Varela and Sherman, 2007, 2009; Hay et al., 2019). Activation of brainstem pedunculopontine area promotes cortical activated states that is prevented by sectioning the thalamus (Steriade et al., 1993).

We show that MTN neurons fire primarily bursts of action potentials during SWS-like and tonic action potentials during REM-like sleep (Figure 2), consistently with previous reports (Zimmerman and Grace, 2018). However, bursting was not suppressed during REM-like state, which contrasts from the reported firing of relay thalamic neurons and could be a feature of higher-order thalamic neurons (Fuster and Alexander, 1973; Weyand et al., 2001; Ramcharan et al., 2005). The proportion of spikes fired in burst during SWS-like was lower than reported by others in the primary sensory thalami but comparable with reports from higher-order nuclei (Zimmerman and Grace, 2018; Urbain et al., 2019). Similarly, burst firing during REM-like state was strikingly high in our sample. In primary sensory thalamic nuclei, burst has been hypothetized to act as a salience detector, and can powerfully activate the cortical primary sensory areas (Swadlow and Gusev, 2001). The role of MTN burst is not yet fully understood. It could allow for maintaining attention during wake (Ramcharan et al., 2005) and during SWS it could promote state transition. Indeed burst firing has been associated with either the initiation of the termination of Up state (Gent et al., 2018; Varela and Wilson, 2020). We did not observe any effect of stimulation or silencing of LDTg neurons on thalamic bursting (Figure 4 and 6). We could extrapolate from these results that at least during sleep-like stage, the cholinergic information relayed by the MTN to the cortex is not meant to promote state transition. Conversely, we observed a reduction of MTN tonic firing when LDTg neurons were silenced and an increase when the cholinergic neurons were stimulated. Tonic firing supposedly acts on the cortical dynamics by promoting state maintenance. During wake, it maintains cortical attention (Schmitt et al., 2017), by promoting the sustained depolarization of the prefrontal cortical neurons. In agreement with these results, the reduced MTN tonic firing in the silencing experiment was associated with shorter Up states (Figure 4). Thus MTN tonic firing could act as a relay for LDTg inputs to regulate cortical arousal state.

### Conclusion

In conclusion, this study aimed at identifying how cholinergic inputs modulate the thalamocortical loop dynamics. It showed that LDTg inputs primarily act by modulating tonic firing in the MTN. As suggested by Peever and Fuller (2016), it is critical to identify how the different structures involved in state regulation communicate with one another. Even though the results presented here do not exclude the possibility that another forebrain relay (for instance the basal forebrain cholinergic neurons) relay brainstem arousal regulating signals to the neocortex, I show evidence that MTN neurons contribute to the dialogue between brainstem arousal centres and the neocortex. It would be essential to continue this work in natural sleep to get a better understanding of the modulation of the thalamocortical loop by neuromodulators.

## ACKNOWLEDGMENTS

I am grateful to Prof Paulsen for his unconditional support, the insightful discussions and to have given me the opportunity to use the resources of his lab to carry this work. This work was supported by a BBSRC grant (BB/S015922/1) and by an Herchel Smith Fellowship. The funder had no role in study design, data collection and analysis, decision to publish, or preparation of the manuscript.

## Notes

**Conflict of Interest:** “The author declares no competing financial interests.”

**Funding:** This work was supported by a BBSRC grant (BB/S015922/1) and by an Herchel Smith Fellowship. The funder had no role in study design, data collection and analysis, decision to publish, or preparation of the manuscript.

### Competing Interest Statement

The authors have declared no competing interest.

## REFERENCES

Bayer L, Eggermann E, Saint-Mleux B, Machard D, Jones BE, Mühlethaler M, Serafin M. Selective action of orexin (hypocretin) on nonspecific thalamocortical projection neurons. J Neurosci. 2002;22(18):7835–9.

Bolkan SS, Stujenske JM, Parnaudeau S, Spellman TJ, Rauffenbart C, Abbas AI, Harris AZ, Gordon JA, Kellendonk C. Thalamic projections sustain prefrontal activity during working memory maintenance. Nat Neurosci. 2017;20:987–96.

Bonjean M, Baker T, Bazhenov M, Cash S, Halgren E, Sejnowski T. Interactions between core and matrix thalamocortical projections in human sleep spindle synchronization. J Neurosci. 2012;32(15):5250–63.

Boucetta S, Jones BE. Activity profiles of cholinergic and intermingled GABAergic and putative glutamatergic neurons in the pontomesencephalic tegmentum of urethane-anesthetized rats. J Neurosci. 2009;29(14):4664–74.

Carracedo LM, Kjeldsen H, Cunnington L, Jenkins A, Schofield I, Cunningham MO, Davies CH, Traub RD, Whittington MA. A neocortical ∂rhythm facilitates reciprocal interlaminar interactions via nested theta rhythms. J Neurosci. 2013;33:10750–61.

Cissé Y, Toossi H, Ishibashi M, Mainville L, Leonard CS, Adamantidis A, Jones BE. Discharge and Role of Acetylcholine Pontomesencephalic Neurons in Cortical Activity and Sleep-Wake States Examined by Optogenetics and Juxtacellular Recording in Mice. eNeuro. 2018;5(4):ENEURO.0270-18.2018.

Cornwall J, Cooper JD, Phillipson OT. Afferent and efferent connections of the laterodorsal tegmental nucleus in the rat. Brain Res Bull. 1990;25:271–84.

Curró Dossi R, Paré D, Steriade M. Short-lasting nicotinic and long-lasting muscarinic depolarizing responses of thalamocortical neurons to stimulation of mesopontine cholinergic nuclei. J Neurophysiol. 1991;65(3):393–406.

David F, Schmiedt JT, Taylor HL, Orban G, Di Giovanni G, Uebele VN, Renger JJ, Lambert RC, Leresche N, Crunelli V. Essential thalamic contribution to slow waves of natural sleep. J Neurosci. 2013;33(50):19599–610.

Dempsey EW, Morison RS. The reproduction of rhythmically recurrent cortical potentials after localized thalamic stimulation. Am J Physiol. 1942;135:293–300.

Eschenko O, Magri C, Panzeri S, Sara SJ. Noradrenergic Neurons of the Locus coeruleus are phase locked to cortical Up-Down states during sleep. Cereb Cortex. 2012;22(2):426–35.

Furman M, Zhan Q, McCafferty C, Lerner BA, Motelow JE, Meng J, Ma C, Buchanan GF, Witten IB, Deisseroth K, Cardin JA, Blumenfeld H. Optogenetic stimulation of cholinergic brainstem neurons during focal limbic seizures: Effects on cortical physiology. Epilepsia. 2015;56(12):e198–202.

Fuster JM, Alexander GE. Firing changes in cells of the nucleus medialis dorsalis associated with delayed response behavior. Brain Res. 1973;61:79–91.

Gent TC, Bandarabadi M, Gutierrez Herrera C, Adamantidis AR. Thalamic dual control of sleep and wakefulness. Nat Neurosci. 2018;21(7):974–84.

Grace KP, Vanstone LE, Horner RL. Endogenous cholinergic input to the pontine REM sleep generator is not required for REM sleep to occur. J Neurosci. 2014;34(43):14198–209.

Guo ZV, Inagaki HK, Daie K, Druckmann S, Gerfen CR, Svoboda K. Maintenance of persistent activity in a frontal thalamocortical loop. Nature. 2017;545:181–6.

Hallanger AE, Levey AI, Lee HJ, Rye DB, Wainer BH. The origins of cholinergic and other subcortical afferents to the thalamus in the rat. J Comp Neurol. 1987;262(1):105–24.

Hauer BE, Pagliardini S, Dickson CT. The Reuniens Nucleus of the Thalamus Has an Essential Role in Coordinating Slow-Wave Activity between Neocortex and Hippocampus. eNeuro. 2019;6(5):ENEURO.0365–19.2019.

Hay YA, Naudé J, Faure P, Lambolez B. Target interneuron preference in thalamocortical pathways determines the temporal structure of cortical responses. Cereb Cortex. 2019;129(7):2815–31.

Hay YA, Jarzebowski P, Zhang Y, Digby R, Brendel V, Paulsen O, Magloire V. Cholinergic modulation of Up-Down states in the mouse medial entorhinal cortex in vitro. Eur J Neurosci. 2021;53(5):1378–93.

Hirsch JC, Fourment A, Marc ME. Sleep-related variations of membrane potential in the lateral geniculate body relay neurons of the cat. Brain Res. 1983;259(2):308–12.

Huerta-Ocampo I, Hacioglu-Bay H, Dautan D, Mena-Segovia J. Distribution of Midbrain Cholinergic Axons in the Thalamus. eNeuro. 2020;7(1):ENEURO.0454-19.2019.

Kitt CA, Höhmann C, Coyle JT, Price DL. Cholinergic innervation of mouse forebrain structures. J Comp Neurol. 1994;341(1):117–29.

Kroeger D, Ferrari LL, Petit G, Mahoney CE, Fuller PM, Arrigoni E, Scammell TE. Cholinergic, glutamatergic, and GABAergic neurons of the pedunculopontine tegmental nucleus have distinct effects on sleep/wake behavior in mice. J Neurosci 2017;37:1352–66.

Lőrincz ML, Gunner D, Bao Y, Connelly WM, Isaac JT, Hughes SW, Crunelli V. A distinct class of slow (∼0.2-2 Hz) intrinsically bursting layer 5 pyramidal neurons determines UP/DOWN state dynamics in the neocortex. J Neurosci. 2015;35:5442–58.

Marrosu F, Portas C, Mascia MS, Casu MA, Fà M, Giagheddu M, Imperato A, Gessa GL. Microdialysis measurement of cortical and hippocampal acetylcholine release during sleep–wake cycle in freely moving cats. Brain Res. 1995;671:329–32.

Mattis J, Tye KM, Ferenczi EA, Ramakrishnan C, O’Shea DJ, Prakash R, Gunaydin LA, Hyun M, Fenno LE, Gradinaru V, Yizhar O, Deisseroth K. Principles for applying optogenetic tools derived from direct comparative analysis of microbial opsins. Nat Methods. 2011;9(2):159–72.

Mena-Segovia J, Sims HM, Magill PJ, Bolam JP. Cholinergic brainstem neurons modulate cortical gamma activity during slow oscillations. J Physiol. 2008;586:2947–60.

Mena-Segovia J, Bolam JP. Rethinking the pedunculopontine nucleus: from cellular organization to function. Neuron. 2017;94(1):7–18.

Mesulam MM, Mufson EJ, Wainer BH, Levey AI. Central cholinergic pathways in the rat: an overview based on an alternative nomenclature (Ch1-Ch6). Neuroscience. 1993;10(4):1185–201.

Moruzzi G, Magoun HW. Brain stem reticular formation and activation of the EEG. Electroencephalogr Clin Neurophysiol. 1949;1(4):455–473.

Pagliardini S, Greer JJ, Funk GD, Dickson CT. State-dependent modulation of breathing in urethane-anesthetized rats. J Neurosci. 2012;32(33):11259–70.

Pagliardini S, Gosgnach S, Dickson CT. Spontaneous sleep-like brain state alternations and breathing characteristics in urethane anesthetized mice. PLoS ONE. 2013;8(7): e70411.

Parent M, Descarries L. Acetylcholine innervation of the adult rat thalamus: distribution and ultrastructural features in dorsolateral geniculate, parafascicular, and reticular thalamic nuclei. J Comp Neurol. 2008;511(5):678–91.

Peever J, Fuller PM. Neuroscience: a distributed neural network controls REM sleep. Curr Biol. 2016;26(1):R34–5.

Petzold A, Valencia M, Pál B, Mena-Segovia J. Decoding brain state transitions in the pedunculopontine nucleus: cooperative phasic and tonic mechanisms. Front Neural Circuits. 2015;9:68.

R Core Team (2018). R: A language and environment for statistical computing. R Foundation for Statistical Computing, Vienna, Austria. URL http://www.R-project.org/

Ramcharan EJ, Gnadt JW, Sherman SM. Higher-order thalamic relays burst more than first-order relays. Proc Natl Acad Sci USA. 2005;102(34):12236–41.

Schmitt LI, Wimmer RD, Nakajima M, Happ M, Mofakham S, Halassa MM. Thalamic amplification of cortical connectivity sustains attentional control. Nature. 2017;545:219–23.

Sheroziya M, Timofeev I. Global intracellular slow-wave dynamics of the thalamocortical system. J Neurosci. 2014;34(26):8875–93.

Steriade M, Nuñez A, Amzica F. A novel slow (< 1 Hz) oscillation of neocortical neurons in vivo: depolarizing and hyperpolarizing components. J Neurosci. 1993a;13(8):3252–65.

Steriade M, Nuñez A, Amzica F. Intracellular analysis of relations between the slow (< 1 Hz) neocortical oscillation and other sleep rhythms of the electroencephalogram. J Neurosci. 1993b;13(8):3266–83.

Steriade M, Contreras D, Curró Dossi R, Nuñez A. The slow (< 1 Hz) oscillation in reticular thalamic and thalamocortical neurons: scenario of sleep rhythm generation in interacting thalamic and neocortical networks. J Neurosci. 1993c;13(8):3284–99.

Steriade M, McCormick DA, Sejnowski TJ. Thalamocortical oscillations in the sleeping and aroused brain. Science. 1993d;262(5134):679–85.

Steriade M, Amzica F, Nuñez A. Cholinergic and noradrenergic modulation of the slow (approximately 0.3 Hz) oscillation in neocortical cells. J Neurophysiol. 1993e;70:1385–1400.

Swadlow HA, Gusev AG. The impact of ’bursting’ thalamic impulses at a neocortical synapse. Nat Neurosci. 2001;4(4):402–8.

Urbain N, Fourcaud-Trocmé N, Laheux S, Salin PA, Gentet LJ. Brain-state-dependent modulation of neuronal firing and membrane potential dynamics in the somatosensory thalamus during natural sleep. Cell Rep. 2019;26(6):1443–1457.e5.

Van Dort CJ, Zachs DP, Kenny JD, Zheng S, Goldblum RR, Gelwan NA, Ramos DM, Nolan MA, Wang K, Weng FJ, Lin Y, Wilson MA, Brown EN. Optogenetic activation of cholinergic neurons in the PPT or LDT induces REM sleep. Proc Natl Acad Sci USA. 2015;112:584–9.

Varela C, Sherman SM. Differences in response to muscarinic activation between first and higher order thalamic relays. J Neurophysiol. 2007;98:3538–3547.

Varela C, Sherman SM. Differences in response to serotonergic activation between first and higher order thalamic nuclei. Cereb Cortex. 2009;19:1776–1786.

Varela C, Wilson MA. mPFC spindle cycles organize sparse thalamic activation and recently active CA1 cells during non-REM sleep. Elife. 2020;9:e48881.

Wester JC, Contreras D. Differential modulation of spontaneous and evoked thalamocortical network activity by acetylcholine level in vitro. J Neurosci. 2013;33(45):17951–66.

Weyand TG, Boudreaux M, Guido W. Burst and tonic response modes in thalamic neurons during sleep and wakefulness. J Neurophysiol. 2001;85(3):1107–18.

Zimmerman EC, Grace AA. Prefrontal cortex modulates firing pattern in the nucleus reuniens of the midline thalamus via distinct corticothalamic pathways. Eur J Neurosci. 2018;48(10):3255–72.

